# Perceptual Decoys Do Not Reliably Bias Choice: Boundary-Condition Evidence

**DOI:** 10.1101/2025.11.02.686051

**Authors:** Daniela Ibarra, Gaurav Suri

## Abstract

The decoy effect occurs when adding an inferior third option biases choice between two others, even though the decoy is rarely chosen. While robust in value-based decisions, evidence in perceptual tasks is mixed. Using the rhesus-macaque paradigm from Parrish et al. (2015), we tested whether a perceptual decoy effect generalizes to humans. Participants (*n* = 50) completed 400 trials. Contrary to our preregistered prediction, we found no reliable decoy effect. Accuracy improved on the hardest trials (Level 1) when a decoy was present, response times were slower in decoy conditions than baseline, and accuracy was higher for tall versus wide rectangles, consistent with the vertical–horizontal asymmetry. The relatively wide spacing of stimuli may have reduced grouping and attentional clustering; because spacing was not manipulated, this remains a hypothesis for future tests. Results suggest that context effects in perceptual choice operate under narrower boundary conditions than in value-based domains.

## Introduction

The decoy effect, used interchangeably with the *asymmetric dominance effect*, is a form of context-dependent bias in which the probability of selecting one of two options is impacted by the addition of a third, inferior alternative (Huber, Payne, & Puto, 1982). The decoy effect occurs when the original two options dominate each other on opposing dimensions. The first option dominates the second on one dimension, while the second option dominates the first on a different dimension. The decoy stimulus is the inferior and irrelevant option in the choice set, yet it serves to “help” the option it most closely resembles by increasing that option’s likelihood of being chosen (Huber et al., 1982).

If a customer walks into an electronics store with hopes to buy a camera that is both economic in price and sufficient in storage, then the dimensions in this scenario are price and storage. If the first camera the customer considers is a $200 camera with 50 GB of storage and the second camera is a $350 camera with 90GB of storage, these two cameras dominate each other on opposing dimensions. The first camera dominates in the dimension of economic price, while the second camera dominates in the dimension of sufficient storage. As the customer walks around the store, another camera catches their attention. This third (*decoy*) camera is $400 and has 80GB of storage. The introduction of this third camera influences the customer to purchase the second camera. While the third camera was inferior to both the original cameras on at least one dimension, it served to increase the probability of selecting the second camera because the decoy was dominated by the second camera on both dimensions. This phenomenon contradicts the rational choice theory (Tversky, 1972), demonstrating how irrelevant alternatives can sway decision-making.

The decoy effect belongs to a family of conceptually related, context-driven phenomena that depend on the availability of other options. These context effects include the attraction or decoy effect (Huber et al., 1982), similarity (Tversky, 1972), and compromise (Simonson, 1989) effects. The extensive research on context effects in value-based tasks finds that decision-making between two options is influenced by the addition of a third alternative, demonstrating how choice behavior is sensitive to contextual factors. Many theoretical models of decision-making that account for these effects, such as the Decision Field Theory (Busemeyer & Townsend, 1993), often highlight attention as a key mechanism driving choice behavior. According to this framework, the decoy effect arises from an attentional bias, where individuals pay more attention to the cluster formed by the decoy and its dominant option than to the standalone alternative.

Research on the decoy effect has been concentrated primarily in the value-based domain, where it has been ubiquitously observed (Doyle, O’Connor, Reynolds, & Bottomley, 1999; Herne, K. 1997; Huber et al., 1982; Kurniawati, Febriyani, & Nicko, 2024; Pan, O’Curry, & Pitts, 1995; Parducci, 1965; Pettibone & Wedell, 2000; Wu, Liu, Chen, Hu, Fan, & Chen, 2020). To investigate the underlying mechanisms of this phenomenon, researchers have sought to replicate the decoy effect in the perceptual domain. Conducting research through perceptual tasks rather than value-based tasks allows for the exploration of context effects at their most basic. There are various advantages to exploring the decoy effect in a perceptual domain. Perceptual tasks elicit quick, low deliberation decisions (Dutilh & Rieskamp, 2016) and minimize the influence of extraneous variables, which are prevalent in value-based decision-making, such as affect and formal calculations of value (e.g., comparing a $200 camera with 50 GB of storage to a $350 camera with 90GB of storage). Perceptual tasks strip away these extraneous influences, making them well-suited for isolating the cognitive mechanisms that drive the decoy effect.

Findings from perceptual studies, however, have been mixed. Some have reported evidence for the decoy effect in perceptual tasks (Choplin & Hummel, 2005; Parrish, Evans, & Beran, 2015; Trueblood, Brown, Heathcote, & Busemeyer, 2013), whereas others have not (Parrish, Dawes, & Thompson, 2024; Spektor, Kellen, & Hotaling, 2018; Spektor, Bhatia, & Gluth, 2021). A strikingly clean design was offered by Parrish et al. (2015), who demonstrated the decoy effect in a perceptual discrimination task with rhesus macaques. In the present study, we sought to extend their paradigm to human participants to determine whether the same effect would emerge. Successful replication of this phenomenon in humans would provide a strong foundation for future investigations into the mechanisms underlying the decoy effect and related context-dependent biases.

It is unknown whether the findings observed in rhesus macaques will replicate in humans. We hypothesize that the decoy effect will emerge. If our hypothesis is on the right track, we anticipate increased accuracy (selection of the largest rectangle) in the congruent condition, relative to the baseline condition. Conversely, we anticipate decreased accuracy in the incongruent condition, relative to the baseline condition. Additionally, we anticipate faster response time in the congruent condition and slower response time in the incongruent condition, relative to the baseline condition.

## Methods

The study pre-registration, programming, data, and code are available on Open Science Framework (OSF): (https://osf.io/6spdn).

### Participants

Fifty undergraduate students from San Francisco State University, all aged 18 or older, participated in the present experiment. Each participant provided verbal and written consent. A priori analysis using G*Power (Faul, F., Erdfelder, E., Lang, A. G., & Buchner, A., 2007) Version 3.1.9.6 was used to determine sample size. We based the expected effect size on the largest within-subjects effect reported in the replicated study (paired-samples *t-*test, *dz* = 4.40). Using this effect size, α = .05, and .80 power, the analysis indicated that a minimum of *N* = 3 was a sufficient number of participants to observe an effect. To ensure stability and generalizability in humans, we recruited a substantially larger sample, far exceeding the minimum requirement. The Institutional Review Board at San Francisco State University approved human participant involvement. Each participant registered through SONA, the university’s psychology research participation system, and received course credit. To incentivize accuracy, the participant with the highest overall accuracy score across all sessions was awarded a $100 Amazon certificate via email after the experiment concluded.

### Apparatus and Stimuli

The task was programmed using PsychoPy Builder (v2024.2.3), which controlled stimulus presentation and data recording. Stimuli were presented on a 24-inch Dell monitor (2408WFPb, Dell Inc., One Dell Way, Round Rock, Texas 78682) at an approximate viewing distance of 120 cm. Participants used a Dell keyboard (Dell L30U Ext. No. D11) and Dell USB laser mouse (MOCZUL DP/N: 0Y357C) on a mousepad to respond during the task.

The stimulus set was modeled after the study by Parrish et al. (2015, Experiment 2), which demonstrated the decoy effect in rhesus macaques (*Macaca mulatta*). All on-screen instructions were presented in black sentence-case font (Open Sans, 0.05 height units). Rectangles appeared in gray on a white background. One rectangle was designated as ‘tall’ (height greater than its width) and the other as ‘wide’ (width greater than its height). In the baseline condition, two rectangles were shown, the largest and the smaller rectangles, randomly positioned on the left or right positions of the screen, with 400 pixels of separation. For half of the trials, the ‘small’ rectangle in the choice set was generated by the program with a width randomly selected between 120 to 169 pixels and a height randomly selected between 60 to 109 pixels. The other half of the trials used a ‘small’ rectangle with a height randomly selected between 120 to 169 pixels and a width randomly selected between 60 to 109 pixels. The ‘large’ rectangle’s height was computed by applying the formula [1 + (Level * 0.015)] to the width of the ‘small’ rectangle. The ‘large’ rectangle’s width was computed by the same formula being applied to the height of the ‘small’ rectangle. This formula, derived from Parrish et al. (2015, Experiment 2), ensures each rectangle appears in both orientations. Level values was randomly generated between 1 to 8 for each trial, with Level 1 being the most difficult (smallest area difference) and Level 8 being the least difficult. The ‘large’ rectangle was equally likely to appear in either orientation. In trials with a ‘decoy’ rectangle (i.e. congruent and incongruent conditions), the ‘decoy’ rectangle had 75% of the area of its same-orientation counterpart. When three rectangles appeared, they were positioned on the left, center, and right of the screen, spaced 400 pixels apart. Rectangle positions and condition types were randomized for each trial.

### Procedure

#### Practice Session

Consistent with the replicated study, Parrish et al. (2015, Experiment 2), each participant began with a practice session to become familiar with selecting the largest rectangle in a choice set. This session included only the baseline condition, where two rectangles, the ‘large’ and ‘small’, were presented in opposing orientations. Participants completed the same 20 baseline trials, with randomized trial order and on-screen positions.

To advance to the test session, participants needed to achieve at least 70% accuracy in the practice session. If this threshold was not met, the session was repeated until the criterion was met. 90% of participants passed the practice session on the first attempt; the remaining 10% succeeded after a second round.

Prior to the practice trials, the participant read a text screen instructing them to select the largest of two rectangles. After clicking a gray rectangle centered on the screen, a 1s blank screen appeared, followed by the presentation of the baseline stimuli. A 1-second blank screen followed between trials. The participant did not receive a visual accuracy score but instead heard a melodic tone for correct responses and a buzz tone for incorrect ones. Participants had up to 10 seconds per trial; failing to respond in time triggered a buzz tone and marked the trial as incomplete. The practice session concluded when the participant pressed the spacebar after completing all 20 trials.

#### Test Session

The test session consisted of 400 trials, divided into three conditions: baseline (70%), congruent (15%), and incongruent (15%). Baseline trials were identical to those completed in the practice session. In the congruent and incongruent conditions, a third, even smaller ‘decoy’ rectangle was introduced to the choice set. In the congruent condition, the decoy matched the orientation of the ‘large’ rectangle; in the incongruent condition, it matched the orientation of the ‘small’ rectangle.

When three rectangles were shown, they were displayed in the left, center, and right positions of the screen, spaced 400 pixels apart. Trial condition and rectangle positioning were randomized. The same formula used during the practice session was applied to compute the rectangle dimensions, with the decoy set to 75% of the area of its same-orientation counterpart. Trial difficulty continued to be randomly selected from Level 1 to 8 across all conditions.

Before the test session began, a text screen instructed the participant to select the largest rectangle from either a two- or three-rectangle set. As in the practice session, after clicking a gray rectangle centered on the screen, a 1s blank screen appeared, followed by the presentation of the trial display. A melodic tone followed correct selections, and a buzz tone followed incorrect ones, each followed by a 1s blank screen before the new trial. The accuracy score was not shown on the screen. The participant continued to have up to 10 seconds to select a rectangle; exceeding the time limit was marked as incomplete. After 400 trials, the session ended when the participant pressed the spacebar.

### Data Analysis

Data analysis was conducted in RStudio (Version 2024.09.0+375) using the *afex* package for repeated-measures ANOVAs and the *stats* package for *t*-tests. Trials with no response (10-s deadline) were marked incomplete and excluded. Trial-level outliers were removed casewise when response time or accuracy values exceeded ±3 SD from a participant’s mean within the relevant analysis. For repeated-measures ANOVAs, participants missing data in any within-subject condition were excluded listwise as required by aov_ez (*afex*). Outlier treatment was not pre-registered.

Accuracy analyses removed 1.28 % of trials overall (Baseline = 1.31 %, Congruent = 0.83 %, Incongruent = 1.57 %). Response time analyses removed 1.97 % overall (Baseline = 1.61 %, Congruent = 2.00 %, Incongruent = 3.60 %). Greenhouse–Geisser corrections were applied when sphericity was violated; multiple comparisons used Bonferroni-adjusted *α*. We report generalized eta squared (*η*^2^_G_) because it is comparable across designs and less biased than partial η^2^ in within-subjects contexts (Bakeman, 2005). Figure 1

**Figure 1.**
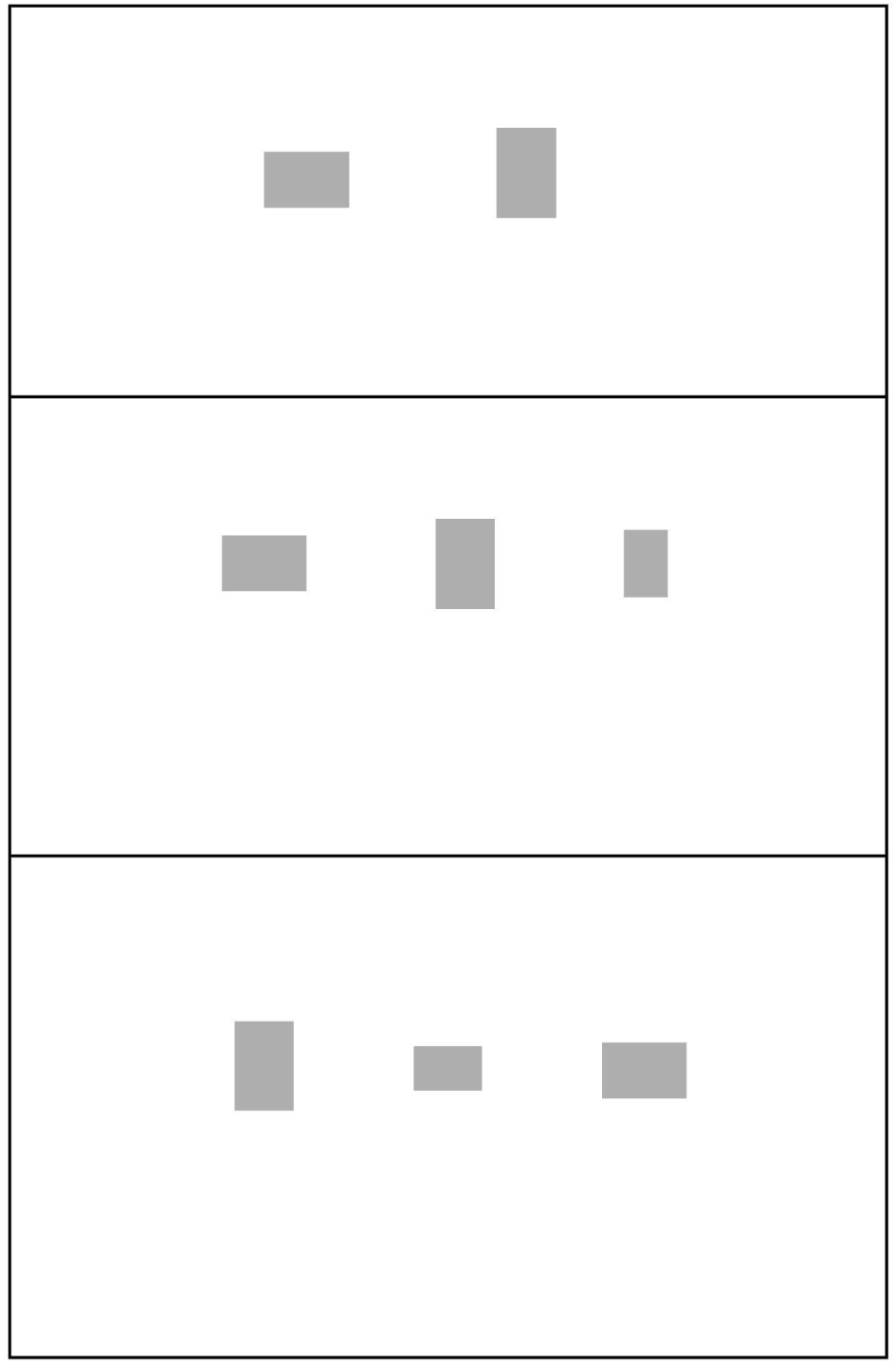
Testing conditions. The top panel shows the baseline condition (no decoy). The middle panel shows the congruent condition (small rectangle on the left, decoy in the middle, and large rectangle on the right). The bottom panel shows the incongruent condition (large rectangle on the left, small rectangle in the middle, and decoy on the right). **Alt text** Visual representation of all three conditions in the testing phase. The top panel shows the baseline condition, the condition where no decoy stimulus is present. In the baseline condition, the small rectangle is on the left and the large rectangle is on the right. The middle panel shows the congruent condition. The small rectangle is on the left, the decoy is in the middle, and the large rectangle is on the right. The bottom panel shows the incongruent condition. Here, the large rectangle on the left, the small rectangle is in the middle, and the decoy is on the right.

## Results

During the testing phase, out of the total 20,000 trials completed (50 participants each completed 400 trials), the decoy rectangle was selected once. This indicates that the decoy rectangle was not a viable choice, as it was rarely (0.005%) selected.

Consistent with the analysis conducted in the replicated study, the data from the eight original difficulty levels were collapsed into four categories by combining pairs of levels. Specifically, Levels 1-2 were combined to form the new Level 1, Levels 3-4 formed the new Level 2, Levels 5-6 formed the new Level 3, and Levels 7-8 formed the new Level 4.

Figure 2 shows the mean percentages of trials correct at each binned stimulus level for each condition and each orientation. Of 50 participants, 14 trials were excluded for no response. Fourteen participant × condition cells (|*z*| > 3) were removed casewise. Eight participant**s** were excluded listwise by *afex* due to incomplete within-subject cells, yielding *n* = 42 for the present analysis. A within-subjects repeated measures analysis of variance (RMANOVA) with a Greenhouse-Geisser correction was conducted to examine the effects of binned level (1–4), condition (baseline, congruent, incongruent), and correct stimulus orientation (tall, wide) on participants’ selection of the larger rectangle. Significant main effects emerged of level *F*(2.60, 106.52) = 184.83, *p* < .001, *η^2^* = .278, and orientation *F*(1, 41) = 33.21, *p* < .001, *η^2^* = .157. However, there was no significant main effect of condition *F*(1.83, 74.86) = 1.15, *p* = .318, *η^2^* = .002, indicating that performance did not vary significantly between the three conditions. There was a significant interaction effect between level and condition, *F*(3.90, 159.92) = 4.36, *p* = .002, *η^2^_G_* = .016, level and orientation, *F*(2.31, 94.64) = 13.58, *p* < .001, *η*^2^_G_ = .042, and between condition and orientation, *F*(1.99, 81.47) = 3.22, *p* = .045, *η^2^_G_* = .005. There was no three-way interaction *F*(4.22, 173.10) = 1.51, *p* = .198, *η*^2^_G_ = .006.

**Figure 2.**
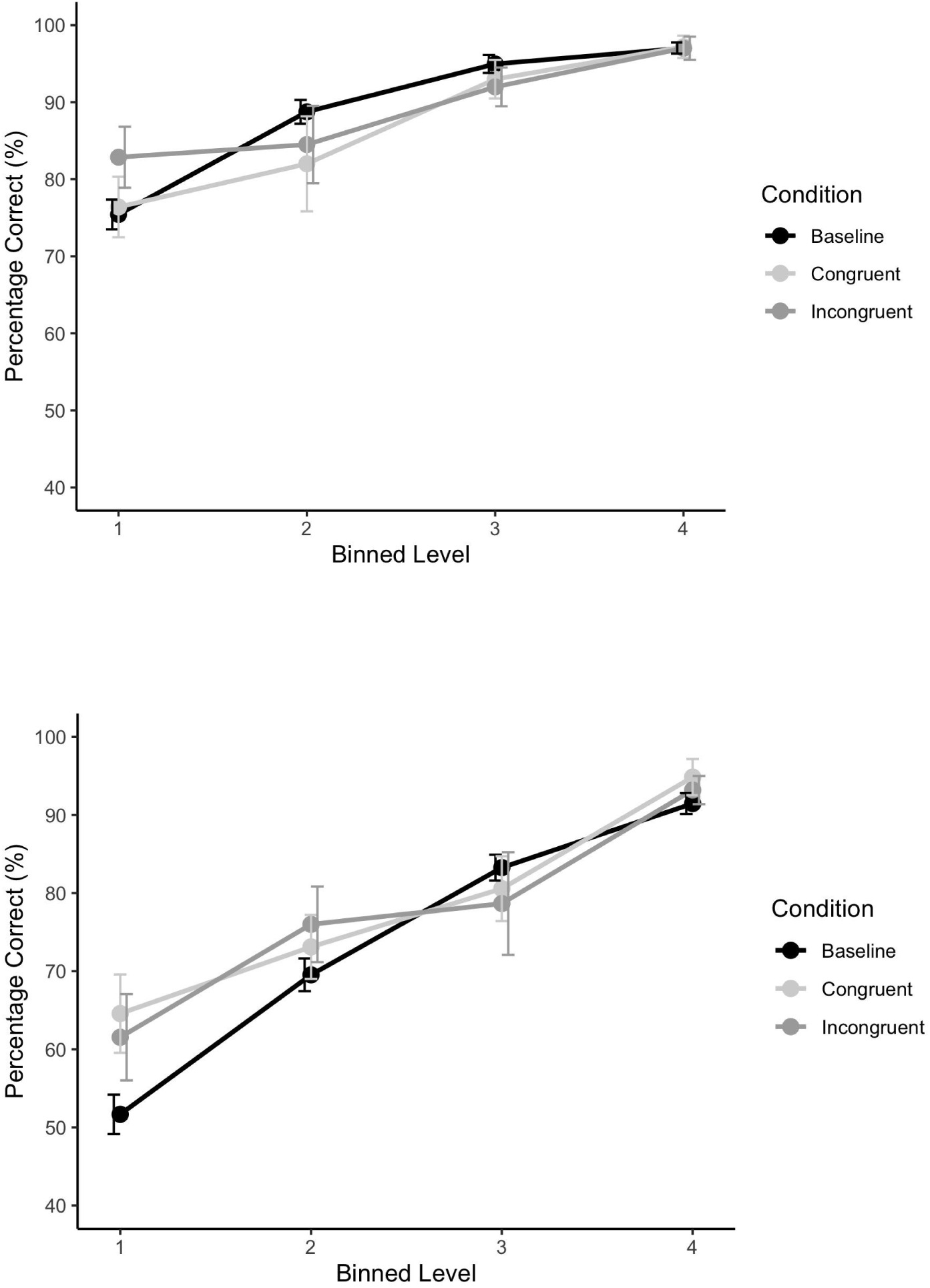
Mean percentage of correct trials at each binned stimulus level (1–4), where Level 1 represents the most difficult discrimination and Level 4 the least difficult, shown separately for each condition (baseline, congruent, and incongruent) and orientation (tall and wide). The top panel depicts the tall orientation, and the bottom panel depicts the wide orientation. Error bars represent 95% confidence intervals. **Alt text:** The top image shows the mean percentage of correct trials at each binned stimulus Level (1–4), with Level 1 representing the most difficult discrimination and Level 4 representing the least difficult discrimination, shown separately for each condition (baseline, congruent, and incongruent), for the tall orientation. The bottom image shows the mean percentage of correct trials at each binned stimulus Level (1–4), with Level 1 representing the most difficult discrimination and Level 4 representing the least difficult discrimination, shown separately for each condition (baseline, congruent, and incongruent), for the wide orientation. Both orientations show a positive slope, demonstrating performance increased at the easier levels. Participants demonstrated higher accuracy for the tall orientation than for the wide orientation. Error bars indicate 95% confidence intervals.

To explore the interaction of level and condition, we compared each condition to each of the others (baseline vs. congruent, baseline vs. incongruent, congruent vs. incongruent) at each level (1, 2, 3, 4) using a paired-samples *t*-test (*n* = 42). A Bonferroni-adjusted *α* level of .017 was used per test (.05/3). Outliers were defined as data points exceeding ±3 SD from the mean difference and were removed casewise where detected: one participant was excluded in Level 2 for the baseline–congruent and congruent–incongruent comparisons, and another participant was excluded in Level 4 for the baseline–incongruent comparison. For Level 1, participants’ performance was significantly higher in the congruent condition (*M* = 0.72, *SD* = 0.11) than in the baseline condition (*M* = 0.66, *SD* = 0.08), *t*(41) = 3.36, *p* = .002, *d* = 0.52. Additionally, participants’ performance was significantly higher in the incongruent condition (*M* = 0.74, *SD* = 0.14) than in the baseline condition (*M* = 0.66, *SD* = 0.08), *t*(41) = 3.35, *p* = .002, *d* = 0.52. Congruent and incongruent conditions were not statistically different, *t*(41) = −0.75, *p* = .456. At Level 2, performance did not differ significantly across conditions [baseline vs. congruent, *t*(40) = 1.56, *p* = .126; baseline vs. incongruent, *t*(41) = −0.24, *p* = .810; and congruent vs. incongruent, *t*(40) = −1.03, *p* = .311]. At Level 3, performance did not differ significantly between conditions [baseline vs. congruent, *t*(41) = 1.10, *p* = .278; baseline and incongruent, *t*(41) = −0.43, *p* = .672; congruent and incongruent, *t*(41) = −1.46, *p* = .153] At Level 4, performance did not differ significantly between conditions [baseline vs. congruent, *t*(41) = −1.59, *p* = .120; baseline vs. incongruent, *t*(40) = −1.32, *p* = .194; congruent vs. incongruent, *t*(41) = 0.86, *p* = .393].

To examine the interaction between orientation and condition, paired-samples *t*-tests were conducted to compare the effects of orientation within each condition, collapsing across levels (*n =* 42). A Bonferroni-adjusted *α* level of .017 (.05/3) was applied. In the baseline condition, performance was significantly higher in tall orientation (*M* = 0.9, *SD* = 0.06) than in wide orientation (*M* = 0.75, *SD* = 0.14), *t*(41) = 5.89, *p* < .001, *d* = 0.91. In the congruent condition, performance was significantly higher in tall orientation (*M* = 0.9, *SD* = 0.08) than in wide orientation (*M* = 0.78, *SD* = 0.13), *t*(41) = 4.19, *p* < .001, *d* = 0.65. In the incongruent condition, performance was significantly higher in tall orientation (*M* = 0.92, *SD* = 0.06) than in wide orientation (*M* = 0.82, *SD* = 0.11), *t*(41) = 4.91, *p* < .001, *d* = 0.76. These results indicate that performance was higher when the correct stimulus appeared in the tall orientation, regardless of condition.

To explore the interaction of orientation and level (this analysis was exploratory as we did not expect this interaction), we compared both orientations to each other (wide vs. tall) at each level (1, 2, 3, 4) using a paired-samples *t*-test (*n* = 42). A Bonferroni-adjusted *α* level of .0125 will be used per test (.05/4). In Level 1, performance was significantly higher in the tall orientation (*M* = 0.79, *SD* = 0.13) than in the wide orientation (*M* = 0.54, *SD* = 0.16), *t*(41) = 6.14, *p* < .001, *d* = 0.95. In Level 2, performance was significantly higher in the tall orientation (*M* = 0.9, *SD* = 0.07) than in the wide orientation (*M* = 0.71, *SD* = 0.17), *t*(41) = 5.87, *p* < .001, *d* = 0.91. In Level 3, performance was significantly higher in the tall orientation (*M* = 0.95, *SD* = 0.05) than in the wide orientation (*M* = 0.83, *SD* = 0.12), *t*(41) = 5.70, *p* < .001, *d* = 0.88. In Level 4, performance was significantly higher in the tall orientation (*M* = 0.98, *SD* = 0.03) than in the wide orientation (*M* = 0.93, *SD* = 0.08), *t*(41) = 4.42, *p* < .001, *d* = 0.68. These results indicate that performance was higher when the correct stimulus appeared in the tall orientation, regardless of difficulty level.

We examined whether response times within a trial differed across conditions for both correct and incorrect trials, based on the assumption that condition effects would be more likely to emerge in correctly completed trials. To test this, a two-way repeated-measures ANOVA with a Greenhouse-Geisser correction was conducted with outcome (correct vs. incorrect) and condition as factors. Mean response times were then calculated for each participant in each condition, separately for correct and incorrect trials. Three participants were excluded listwise (*afex* listwise criterion), leaving *n* = 47. There were significant main effects of condition, *F*(1.89, 86.97) = 18.61, *p* < .001, *η^2^* = .032, and outcome, *F*(1, 46) = 53.96, *p* < .001, *η*^2^ = .072, on response times. However, there was no significant interaction between condition and outcome *F*(1.97, 90.44) = 1.45, *p* = .241, *η*^2^_G_ = .002.

We conducted separate repeated measures ANOVAs with a Greenhouse-Geisser correction for correct and incorrect trials, comparing the response times in each condition (*n* = 47; Fig. 3). We found significant effects of condition for correct trials *F*(1.74, 80.01) = 55.21, *p* < .001, η^2^ = .067, and incorrect trials, *F*(1.90, 87.17) = 4.93, *p* = .010, *η*^2^ = .020.

**Figure 3.**
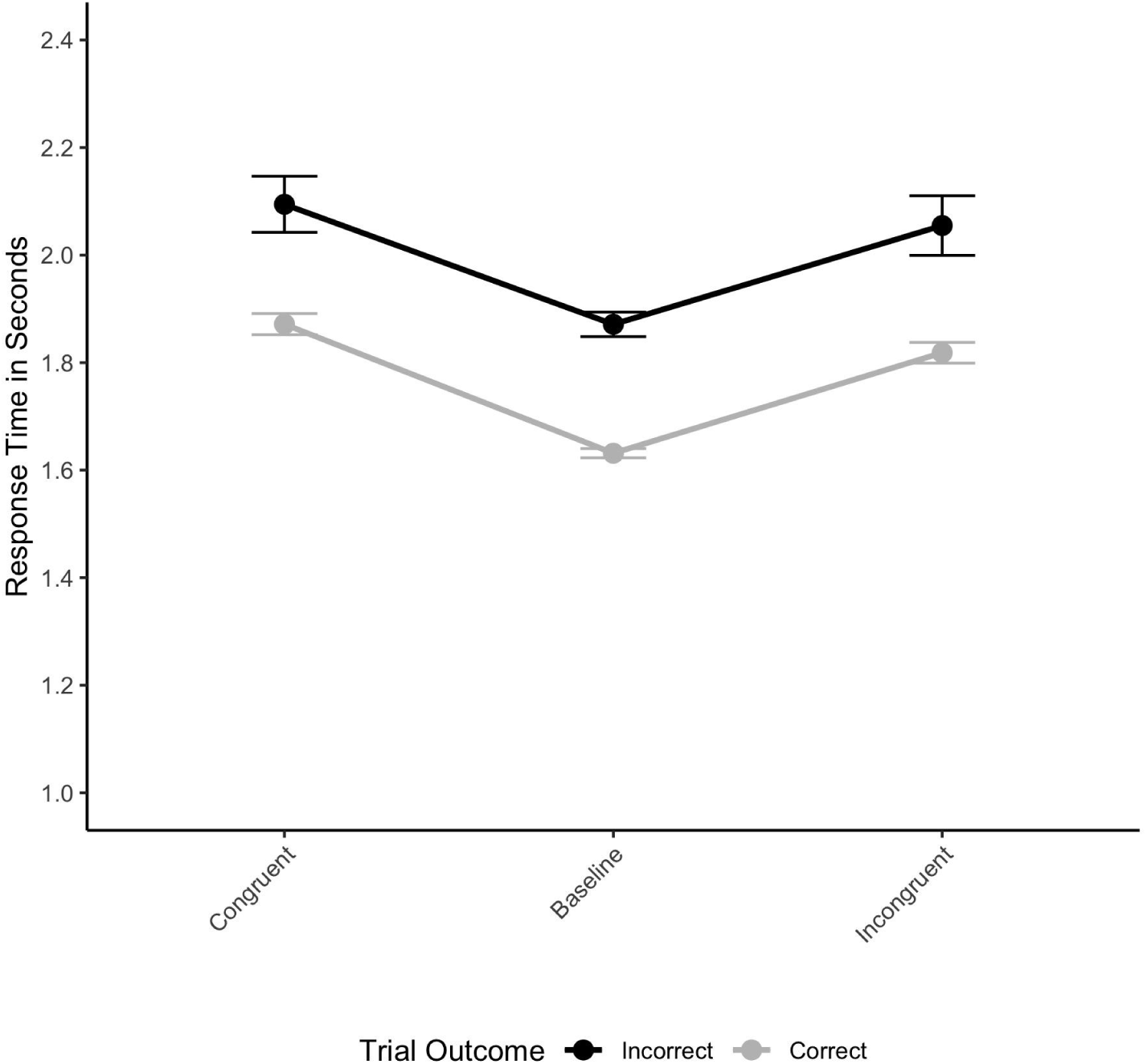
Mean response times by condition for correct and incorrect trials. Error bars represent standard errors of the mean. **Alt text:** The figure shows the mean response times by condition for correct and incorrect trials. Across both correct and incorrect trials, response times were longer for the congruent and incongruent conditions than in the baseline condition. Error bars represent standard errors of the mean.

We then compared each condition to the others using paired-samples *t*-tests for correct and incorrect trials (*n* = 47; Bonferroni-adjusted *α =* .017). Outliers were defined as data points exceeding ±3 SD from the mean difference and were removed casewise when detected. For correct trials, two participants were excluded for the baseline–congruent comparison, and one participant was excluded for the incongruent–congruent comparison. In correct trials, participants’ response times were significantly faster in the baseline condition (*M* = 1.55, *SD* = 0.33) than in the congruent condition (*M* = 1.76, *SD* = 0.39), *t*(44) = 10.74, *p* < .001, *d* = 1.60. Additionally, participants’ response times were significantly faster in the baseline condition (*M* = 1.57, *SD* = 0.33) than in the incongruent condition (*M* = 1.74, *SD* = 0.38), *t*(46) = 9.53, *p* < .001, *d* = 1.39. There was no significant effect between participants’ response times in the incongruent condition (*M* = 1.74, *SD* = 0.38) and the congruent condition (*M* = 1.79, *SD* = 0.42), *t*(45) = 2.22, *p* = 0.032, *d* = .33. In incorrect trials, participants’ response times were significantly faster in the baseline condition (*M* = 1.87, *SD* = 0.58) than in the congruent condition (*M* = 2.02, *SD* = 0.61), *t*(46) = 2.54, *p* = 0.015, *d* = 0.37. Additionally, participants’ response times were significantly faster in the baseline condition (*M* = 1.87, *SD* = 0.58) than in the incongruent condition (*M* = 2.08, *SD* = 0.66), *t*(46) = 2.87, *p* = 0.006, *d* = 0.42. There was no significant difference between participants’ response times in the incongruent condition (*M* = 2.08, *SD* = 0.66) and the congruent condition (*M* = 2.02, *SD* = 0.61), *t*(46) = 0.77, *p* = 0.443, *d* = .11.

## Discussion

The present study aimed to determine whether the decoy effect, a phenomenon ubiquitously observed in value-based decision-making, would extend to perceptual choice. Our findings show that under conditions mirroring the Parrish et al. (2015) paradigm, we found no reliable evidence of a decoy effect, suggesting that the mechanisms underlying perceptual context effects may differ from those driving value-based context effects. Consistent with this interpretation, our findings reveal minimal influence of the decoy itself. Participants rarely selected the decoy, and accuracy did not differ overall across baseline, congruent, and incongruent conditions. Instead, three main results emerged: (a) accuracy improvement on the most difficult trials when a decoy was present, (b) slower response times in decoy conditions compared to baseline, and (c) a robust orientation effect favoring tall rectangles, consistent with the vertical–horizontal illusion (Künnapas, 1955; Mamassian & de Montalembert, 2010). These findings suggest that perceptual asymmetries shaped performance and that adding a third item increased processing demands without reliably biasing choice. Together, the null on choice, the reaction time cost of adding an alternative, and the tall–wide asymmetry point to perceptual constraints and attentional allocation as boundary conditions for attraction effects in perceptual tasks.

Our findings contribute to a growing body of literature reporting null results for the decoy effect in the perceptual domain. This contrast is striking given that the decoy effect is ubiquitous in the value-based domain. This discrepancy highlights an imperative question: What is the fundamental difference between perceptual and value-based decision processes?

Trueblood et al. (2013) argued that context effects are a fundamental feature of decision-making processes, a position with which we agree. We extend this perspective by proposing that value itself is a context variable, one that is so attentionally salient that it may engage in formal systems and, in doing so, overwhelms or washes away other contextual influences. Consequently, the mechanisms underlying context effects may differ between perceptual and value-based domains. In value-based tasks, affective evaluation, high attention salience, and formal logic reasoning about value calculations all contribute to decision outcomes.

In contrast, perceptual tasks require attention to stimulus-level and design-related factors. In the perceptual domain, the literature includes null findings and even reversals under manipulations such as spacing, attribute distance, grouping, and time pressure (Pettibone, J.C. 2012; Spektor, Kellen, & Hotaling, 2018; Spektor, Bhatia, & Gluth, 2021). One possible explanation for the absence of a decoy effect is that the relatively wide spatial separation between stimuli may have reduced opportunities for grouping and attentional clustering. However, because spacing was not experimentally manipulated, this interpretation remains speculative and should be tested in future research. This interpretation aligns with models emphasizing the role of grouping, similarity, and attentional allocation in context effects (Izakson, Zeevi, & Levy, 2020; Spektor et al., 2021). In particular, Gestalt principles such as proximity may be critical for enabling perceptual decoy-driven biases. When grouping is disrupted, as in our design with wide spacing, the decoy may remain perceptually isolated, leading to slower processing but no systematic bias in choice.

Our findings provide boundary-condition evidence suggesting that perceptual and value-based decision processes may rely on partly distinct mechanisms. However, stronger claims will require direct cross-domain comparisons and convergent evidence across paradigms. Value may act as a dominating contextual variable that minimizes other influences in value-based domains, whereas perceptual context effects may rely on design or display features of the stimuli. Any mechanistic theory of the decoy effect, and of any other related context-dependent bias, must therefore take the saliency of context across both domains into account.

## Contributions

The article has been reviewed and edited by ChatGPT-5, and ChatGPT-5 was also used for coding in RStudio as part of the formal analysis.

Conceptualization: GS, DI

Data Curation: DI

Formal Analysis: DI, AI

Funding Acquisition: N/A

Investigation: DI, GS

Methodology: DI, GS

Project Administration: DI, GS

Resources: DI

Software: DI

Supervision: GS, DI

Validation: GS, DI

Visualization: DI

Writing - original draft: DI

Writing - review and editing: DI, GS, AI

## Acknowledgements

The authors would like to thank the research assistants in the Readiness, Activation, and Decision-making Laboratory at San Francisco State University who assisted with data collection.

## Social Impact and Responsibility

This study investigates perceptual choice behavior and boundary conditions of the decoy effect, contributing to a more robust understanding of context-dependent decision-making. The findings have limited social impact but contribute to scientific integrity by documenting null results.

## Ethics Statement

IRB approval for this study was obtained (2024-582) by San Francisco State University.

## Data Accessibility

The present research was pre-registered on Open Science Framework (OSF): https://osf.io/6spdn/overview

Additionally, the programming, data, and analysis codes used have been made public on OSF: https://osf.io/qwkx6/overview

The repository includes a file named *“Reproducibility Statement”* describing the analysis software, necessary packages, and instructions for reproducing the reported analysis.

